# Dynamic Change of Electrostatic Field in TMEM16F Permeation Pathway Shifts Its Ion Selectivity

**DOI:** 10.1101/515569

**Authors:** Wenlei Ye, Tina W. Han, Mu He, Yuh Nung Jan, Lily Y. Jan

## Abstract

TMEM16F is activated by elevated intracellular Ca^2+^, and functions both as a small-conductance ion channel permeable to Ca^2+^ and as a phospholipid scramblase. It is important to hold this positive feedback in check to prevent prolonged Ca^2+^-overloading in cells. We hypothesize that TMEM16F shifts its ion selectivity so that it is more permeable to Cl^−^ than cations at high intracellular Ca^2+^ concentration. We tested this hypothesis with the Q559K mutant that shows no current rundown in excised patch with prolonged Ca^2+^ elevation. Recorded in NaCl^−^based solution, the channel shifted its ion selectivity from Na^+^-selective to Cl^−^-selective when intracellular Ca^2+^ was increased. The ion selectivity switch did not correlate with changes of channel open state. Rather, it was indicative of an alteration of electrostatic field in the permeation pathway. Shifting ion-selectivity synergistically by intracellular divalent ions and membrane potential could work as a built-in mechanism that allows TMEM16F to brake the positive feedback.

## Introduction

Mammalian TMEM16F is a membrane protein with dual functions of phospholipid scrambling and ion conduction, both activated by elevation of intracellular Ca^2+^ (1–4). When activated, TMEM16F mediates the exposure of phosphatidylserine, a lipid normally restricted to the inner leaflet of cell membrane lipid bilayer, to the cell surface (5). This process, known as lipid scrambling, initiates many physiological processes such as recruitment of thrombin to the platelet surface to trigger blood coagulation and induction of immune responses in T lymphocytes (5–8). TMEM16F is also a Ca^2+^-activated small-conductance ion channel, yielding non-selective cation current in excised patch recordings (6, 9). Consistently, activation of TMEM16F increases cytoplasmic Ca^2+^ concentration in cells that heterologously express this channel (10). The intracellular Ca^2+^ elevation might further activate TMEM16F, forming positive feedback that could potentially lead to detrimental consequences. We have shown that in response to high cytoplasmic Ca^2+^, degradation of phosphatidylinositol-(4,5)-bisphosphate (PIP_2_), a lipid mostly in the cytoplasmic leaflet of cell membrane, could serve as a brake to terminate TMEM16F activation (11). For this brake mechanism to be effective, PIP_2_ degradation needs to be faster than its synthesis in native cells. Moreover, the residual TMEM16F current may still be able to contribute to the positive feedback. We hence searched for intrinsic mechanisms to terminate the positive feedback resulting from TMEM16F activation by Ca^2+^ and its ability to permeate Ca^2+^ ions.

There is discrepancy regarding TMEM16F ion selectivity reported by different labs. Recorded with excised-patch inside-out configuration, TMEM16F channels are quickly activated by micromolar Ca^2+^ and they are more permeable to cations (mainly physiological cations such as Na^+^, K^+^ and Ca^2+^) than Cl^−^, while prolonged treatment by Ca^2+^ leads to current inactivation (6, 9). Surprisingly, many groups reported that TMEM16F whole-cell current is activated several minutes after Ca^2+^ elevation, and it displays less cation-selectivity (12) or even greater selectivity for Cl^−^ than Na^+^ (13–16). This paradox has not been resolved, but nonetheless it suggests that prolonged treatment with high cytoplasmic Ca^2+^ is correlated with a reduction of TMEM16F cation selectivity. Interestingly, recent studies suggested that intracellular Ca^2+^ might alter the accessibility of different ions to the permeation pathway of TMEM16A, a Ca^2+^-activated anion channel paralog of TMEM16F (17). We thus hypothesize that TMEM16F shifts its ion selectivity in response to elevation of intracellular Ca^2+^ concentrations. However, because TMEM16F current in inside-out excised patch exhibits rapid desensitization and rundown in high Ca^2+^, it is challenging to test for TMEM16F ion selectivity under such a variety of Ca^2+^ concentrations.

Here, we report that TMEM16F Q559K current persisted with prolonged exposure to high intracellular Ca^2+^. Previous studies have indicated that Q559 faces the ionic permeation pathway and that lysine substitution reduces TMEM16F cation selectivity (6, 9), and we now further show that this mutant displays different Na^+^/Cl^−^ selectivity in different Ca^2+^ concentrations. The difference in ion selectivity did not correlate with alterations of the open states, but instead was regulated by the electrostatic change along the permeation pathway, on which divalent ions (such as Ca^2+^ and Zn^2+^) had more significant impact than monovalent ions. Furthermore, membrane depolarization, which facilitates intracellular cation entry into the membrane electric field, promoted the selectivity shift toward a preference for Cl^−^ over cations. The shift of cation/anion selectivity could reflect a general feature of how cytoplasmic ions might influence the function of TMEM16 proteins, and it suggests that TMEM16F harbors an inherent machinery that allows it to switch from being cation-selective to being Cl^−^-selective to brake the signals and the ensuing positive feedback.

## Results

### TMEM16F Q559K shifts ion selectivity towards Cl^−^ as intracellular Ca^2+^ increases

Recorded with inside-out configuration and held at +80 mV, wild-type TMEM16F current was activated by intracellular Ca^2+^ in a dose-dependent manner. The current started to decrease when Ca^2+^ was higher than ~30 μM, as a result of both desensitization and rundown (Figure 1A, C, D) (11). Here, desensitization refers to a decreased sensitivity to Ca^2+^, caused by degradation of PIP_2_ via membrane-tethered phospholipase activated by high intracellular Ca^2+^. Rundown refers to a reduction of fully-activated current magnitude (induced by 1 mM Ca^2+^ in this case). The rapid decrease of current rendered it difficult to record the reversal potential of wild-type TMEM16F in 1 mM Ca^2+^, because when we switched the bath (equivalent to intracellular solution) from 150 mM NaCl to 15 mM NaCl, the currents recorded from Tmem16f-transfected cells were indistinguishable from currents endogenous to HEK293 cells (Figure 1–figure supplement 1A, B, C). Consistent with previous reports (6, 9), TMEM16F current induced by 15 μM Ca^2+^ was selective for Na^+^ over Cl^−^ (Figure 1E).

**Figure 1.**
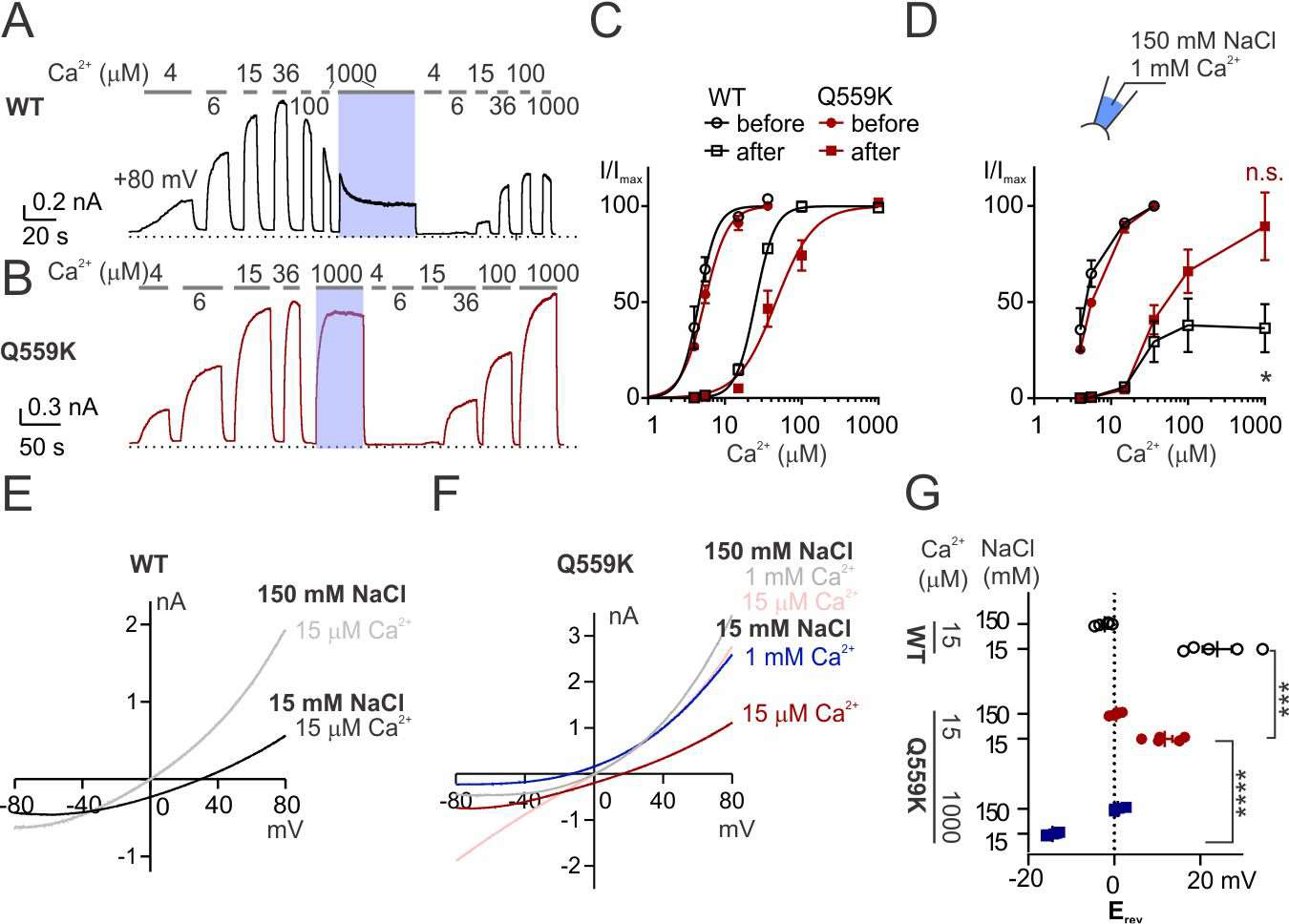
TMEM16F Q559K shifts its reversal potential in response to change of intracellular Ca^2+^ concentration. (*A, B*) Representative recordings of TMEM16F wild type (WT) and Q559K in different Ca^2+^ concentrations. Traces were recorded from transfected HEK cells and the inside-out patches were held at +80 mV. The shades illustrate 1-minute treatment with 1 mM Ca^2+^ that catalyzes PIP_2_ degradation by membrane-tethered phospholipase. (*C*) Dose-response curves for Ca^2+^-activation of WT and Q559K before and after 1 mM Ca^2+^ treatment respectively. Currents before and after 1mM Ca^2+^ were separately fitted to Hill equation and normalized to their respective maximal amplitudes. (*D*) Change of current magnitudes of WT and Q559K. The currents were normalized to the maximal magnitudes before 1 mM Ca^2+^ for each cell. * P < 0.05 (One sample t test against hypothetical value “1”). (*E, F*) Representative I-V relationships of WT and Q559K recorded in indicated conditions. The traces were recorded with a hyperpolarizing ramp from +80 mV to −80 mV (−1 V/s) following holding at +80 mV. (*G*) Scatter plot of reversal potentials (E_rev_) obtained from traces as in *E* and *F. P* values were determined with Sidak’s multiple comparisons following two-way ANOVA

In contrast, the mutant Q559K current recorded in the same condition showed minimal rundown in 1 mM Ca^2+^, in spite of desensitization to Ca^2+^-activation likely due to PIP_2_ depletion. Normalized to the maximal magnitudes of the current before and after 1-minute-treatment of 1 mM Ca^2+^, EC_50_ of Ca^2+^-activation was shifted from 5.6 ± 0.3 μM to 52 ± 11 μM (Figure 1B, C), while the normalized fully-activated current magnitude was 0.89 ± 0.18 (current in 1 mM Ca^2+^ normalized to that before 1 mM Ca^2+^), significantly different from 0.36 ± 0.13 for wild type (Figure 1D). We recorded with a voltage-family protocol from −40 mV to +160 mV with 10 mV increment followed by holding at 100 mV to obtain “tail-currents”, which we used as indicators of conductance. Fitting to the sigmoidal G-V relationship yielded V_1/2_ of 76 ± 11 mV in 15 μM Ca^2+^ for wild-type TMEM16F, while in 1 mM Ca^2+^, the depletion of PIP_2_ strongly reduced the voltage-gating, consistent with our previous reports (Figure 1–figure supplement 2A, D) (11). Depletion of PIP_2_ caused a right-shift of the voltage-dependence of Q559K current in 15 μM Ca^2+^, but in 1 mM Ca^2+^ the left-shift otherwise overrode the effect of PIP_2_ depletion, with V_1/2_ being 18 ± 10 (Figure 1–figure supplement 2B, C, D). Thus, the mutation of Q559K allowed us to record TMEM16F current around physiological membrane voltages in a wide-range of Ca^2+^, here particularly, to measure E_rev_ in 1 mM Ca^2+^.

Interestingly, the E_rev_ of TMEM16F Q559K current activated by 1 mM Ca^2+^ exhibited a left-shift to −14 ± 1 mV in 15 mM NaCl, suggesting that it was selective for Cl^−^ over Na^+^ (Figure 1F, G and Figure 1–figure supplement 1D). Corrected for liquid junction potentials calculated with Clampex, this result indicated that the permeability ratio for Na^+^ over Cl^−^ was 0.47 ± 0.03, in contrast to 2.1 ± 0.4, calculated from Erev in 15 μM Ca^2+^. To test whether the inversion of ion selectivity was due to PIP_2_ depletion, we compared the values of Erev in 15 μM Ca^2+^ before and after perfusion of poly-L-lysine (PLL), which could sequester PIP_2_ from the membrane. After PLL treatment the current in 15 μM Ca^2+^ recorded at +80 mV was too small, so for this experiment we held the patch at +160 mV followed by the hyperpolarizing ramp. Judging from the currents with acceptable magnitudes, the Erev shift was minor, significantly different from that in 1 mM Ca^2+^ (Figure 1–figure supplement 1E, F). Thus, PIP_2_ depletion cannot account for the inversion of ion selectivity of Q559K current.

TMEM16F functions as a cation channel because it is more permeable to physiological cations (Na^+^, K^+^ and Ca^2+^) than Cl^−^, even though the permeability for many other cations may be lower than that for Cl^−^ (6, 12). Like wild-type TMEM16F, Q559K channels are more permeable to I− than Cl^−^ (Figure 1–figure supplement 3). In this study, we will mainly focus on the comparison between Na^+^ permeability and Cl^−^ permeability. Given the variable P_Na+_/P_Cl−_, we cannot employ the calculation methods normally used for bi-ionic conditions to analyze data obtained with solutions involving a third ion species.

### Q559K channel ion selectivity varies with intracellular Ca^2+^ concentration

We asked whether the two phases of ion selectivity of the Q559K mutant channel corresponds to multiple open states that may correlate with different extent of occupancy of the Ca^2+^ binding sites; For example, the channel may be in an intermediate state in 15 μM Ca^2+^ but a fully open state in 1 mM Ca^2+^. To test this possibility, we made use of the Ca^2+^-binding-site mutant, E667Q, whose EC50 of Ca^2+^-activation is shifted to the right, to dissociate the effect of Ca^2+^ concentration from those attributable to different open states of the channel (6, 9). The EC50 of the double-mutant, Q559K_E667Q, was 0.88 ± 0.06 mM, suggesting that the double mutant should not be in the fully open state in 1 mM Ca^2+^ (Figure 2A, B). However, the Erev of the current activated by 1 mM Ca^2+^ was −6.9 ± 2.0 mV, corresponding to a Cl^−^ selective channel (Figure 2C, D). This indicates either that TMEM16F ion selectivity is not correlated with its open state, or that the E667Q mutation circumvents the intermediate open state (if any) and causes the mutant channel to directly enter a fully-open conformation.

**Figure 2.**
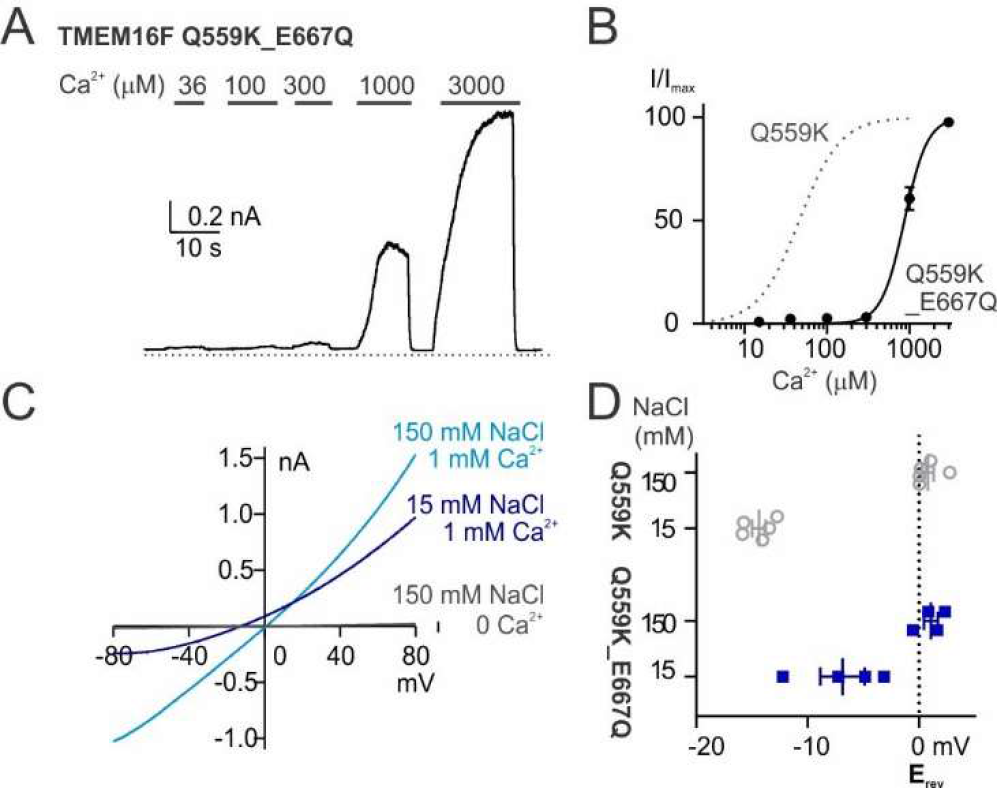
TMEM16F Q559K_E667Q current is Cl^−^ -selective in 1 mM Ca^2+^ despite being half activated. (A) Representative recordings of TMEM16F Q559K_E667Q in different Ca^2+^ concentrations. The recording protocol was the same as in Figure 1A. (*B*) Dose-response curves for Ca^2+^-activation of Q559K_E667Q. The gray dotted line represents the curve for Q559K after 1 mM Ca^2+^, replotted from Figure 1C. (*C*) Representative I-V relationships of Q559K_E667Q recorded in indicated conditions. The recording protocol was the same as in Figure 1E. (*D*) Scatter plot of reversal potentials (E_rev_) obtained from traces as in *C*. The gray values were replotted from Figure 1G for comparison

To further investigate the activation of TMEM16F, we aligned the sequence of its transmembrane helix 6 (TM6) with that of TMEM16A, which plays a critical role in TMEM16A channel gating. In TMEM16A, the glycine in TM6 (G640 or G644 depending on isoform) works as a hinge to allow the rearrangement of the helical segments during channel activation to generate an ionic permeation pathway (18, 19); Alanine substitution of this glycine stabilizes TM6 at the open conformation, and increases the Ca^2+^ sensitivity of TMEM16A (18, 19). Similarly, alanine substitution of G615 of TMEM16F increased the Ca^2+^ sensitivity (9, 10) and caused a left-shift of the voltage dependence recorded in 15 μM Ca^2+^ (Figure 3A, B). Notably, the glutamine at the lower segment of TM6 in TMEM16A, Q645, whose alanine substitution stabilizes the channel at a conformation similar to wild-type TMEM16A bound to one Ca^2+^ ion (18), corresponds to a gap of the alignment in TMEM16F (Figure 3–figure supplement 1A). Alanine substitution of the asparagine “5-amino-acid-away” in TMEM16F TM6, N620, did not alter Ca^2+^ sensitivity (10) or voltage gating (Figure 3–figure supplement 1B, C). In contrast, alanine substitution of the isoleucine I612 caused shifts in Ca^2+^− and voltage-gating (10) comparable to the effect of alanine substitution of I637 at the corresponding position in TMEM16A (Figure 3–figure supplement 1D, G) (18). The comparison of gating between TMEM16A and TMEM16F is beyond the scope of this manuscript but will be briefly discussed later.

**Figure 3.**
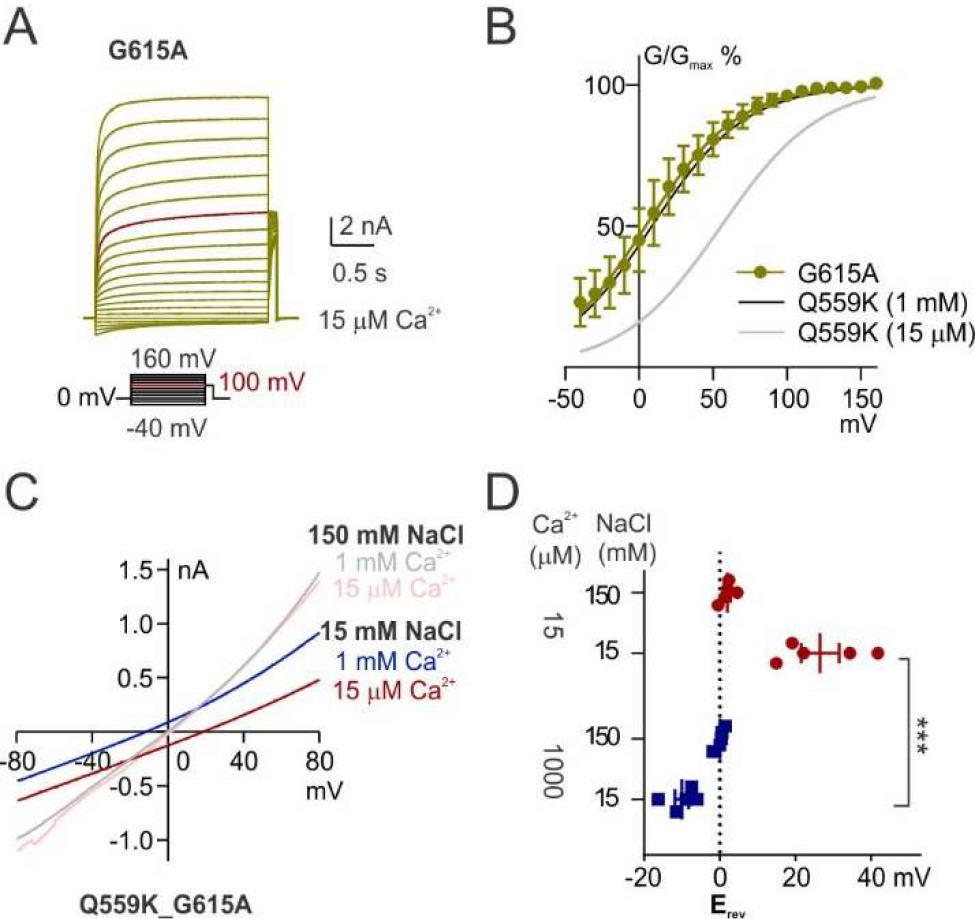
The shift of ion selectivity is preserved despite TM6 conformational fixation. (*A*) Representative trace of G615A recorded with a voltage family protocol. The current recorded at +100 mV is highlighted for comparison. (*B*) Averaged G-V relationships of G615A currents. The method for data analysis was the same as that for WT in 15 μM Ca^2+^. The two traces for Q559K were replotted from Figure 1_Supplement 2D. (*C*) Representative I-V relationships of Q559K_G615A recorded in indicated conditions. (*D*) Scatter plot of reversal potentials (E_rev_) obtained from traces as in *C*. *P* values were determined with Sidak’s multiple comparisons following two-way ANOVA

Notably, the voltage-dependence of TMEM16F G615A channels in 15 μM Ca^2+^ was comparable to that of Q559K channels in 1 mM Ca^2+^ (Figure 3A, B), suggesting that removing the TM6 glycine hinge also stabilized the TMEM16F open conformation. We then generated the double-mutant Q559K_G615A to study its ion selectivity. The switch from Na^+^-selective to Cl^−^-selective was preserved in this double mutant (Figure 3C, D). Our study of two different double mutants reveal that, in the presence of Q559K, 15 μM Ca^2+^ elicited a cation-selective current regardless of TM6 stabilization (Figure 3), while current in 1 mM Ca^2+^ was Cl^−^-selective even if the channel was not fully activated to reach the plateau level in the Ca^2+^ dependence curve (Figure 2). These results suggest that TMEM16F ion selectivity more likely depends on intracellular Ca^2+^ concentration directly rather than its open state(s), and it remains possible that the selectivity could be influenced by other regulatory factors.

### The switch of ion selectivity correlates with electrostatic change in permeating pathway

Intrigued by the possibility that Ca^2+^ alters anion accessibility to TMEM16A pore through electrostatic effect (17), we hypothesize that the shift of ion selectivity of TMEM16F Q559K in response to elevation of intracellular Ca^2+^ is due to a change of electrostatic field along its permeation pathway. In this scenario, the effect ought to be elicited not only by Ca^2+^ but also by other divalent cations (17). Here, we chose to test Zn^2+^, a divalent cation that is smaller but also capable of activating TMEM16F. Wild-type TMEM16F activation by 1 μM Zn^2+^ was similar in magnitude to that by ~10 μM Ca^2+^ with or without PIP_2_ depletion (Figure 4A, B). Zn^2+^-activation of TMEM16F was coupled with rapid inactivation, precluding the possibility to accurately plot the dose-response curve. Since exposure to divalent-free solution (with EGTA) for ~30 s to 1 min following channel activation by Zn^2+^ allowed the TMEM16F current, whether activated by Ca^2+^ or by Zn^2+^, to fully recover in magnitude (Figure 4–figure supplement 1), the rapid current rundown cannot be attributed to desensitization triggered by degradation of PIP_2_.

**Figure 4.**
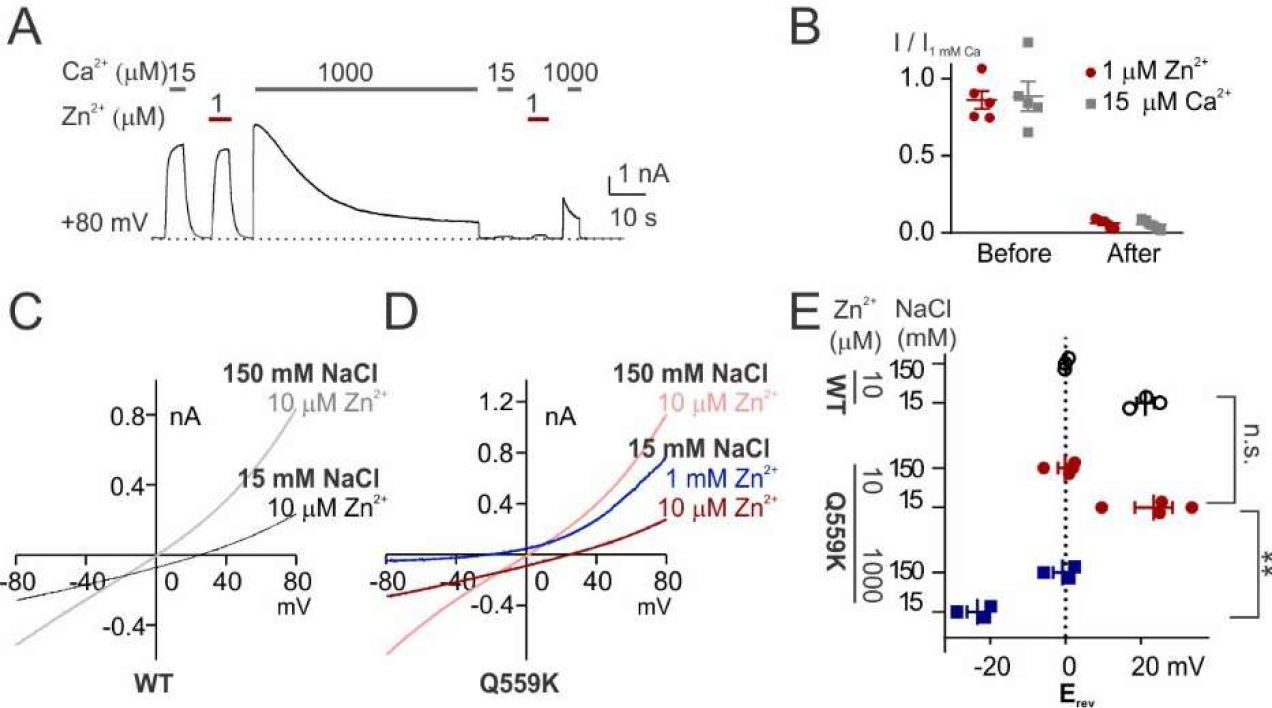
The shift of ion selectivity is preserved when current is activated by Zn^2+^. (*A*) Representative recordings of WT TMEM16F current in response to indicated concentrations of Zn^2+^ and Ca^2+^. (*B*) Scatter plot showing the current magnitudes activated by indicated concentrations of Zn^2+^ and Ca^2+^ normalized to those by 1 mM Ca^2+^ respectively before and after prolonged treatment of 1 mM Ca^2+^, suggesting that Zn^2+^ activation was also desensitized. Two-way ANOVA suggests there is no significant difference between the two activators, but there is significant difference before and after 1 mM Ca^2+^. (*C, D*) Representative I-V relationships of WT and Q559K recorded in indicated conditions. The currents were recorded with the same protocol as in Figure 1E, F. (*E*) Scatter plot of reversal potentials (E_rev_) obtained from traces as in *C* and *D*. *P* values were determined with Sidak’s multiple comparisons following two-way ANOVA

We found that the wild-type TMEM16F channels activated by 10 μM Zn^2+^ was selective for Na^+^ over Cl^−^, as evidenced from the right-shift in E_rev_ as intracellular NaCl dropped to 15 mM (Figure 4C). The reversal potential in 1 mM Zn^2+^ could not be determined owing to the rapid current inactivation. TMEM16F Q559K channels also exhibited a shift from being Na^+^-selective to being Cl^−^-selective with elevation of Zn^2+^ concentration from 10 μM to 1 mM (Figure 4C, D, E). The shift of ion selectivity therefore represents an inherent feature of TMEM16F Q559K channels regardless of the divalent ion species.

Although monovalent ions generate weak electrostatic fields compared with divalent ions, due to their abundance they should also be able to regulate the electrostatic field along TMEM16F permeation pathway and shift the ion selectivity. TMEM16F is permeable to most generally-used cations (6, 12), so introducing any other ion will cause difficulties in distinguishing whether the shift of E_rev_ is due to the change of P_Na+_/P_Cl−_ or to the permeation of the third ion species. Thus, we measured the E_rev_ with different intracellular NaCl concentrations and calculated the respective permeability ratios. When bath (equivalent to intracellular solution) was changed from 15 mM NaCl to 45 mM NaCl and the holding potential was held constant at +80 mV, the driving potential for Na^+^ moving across the membrane was changed by more than 2 folds (from ~20 mV to ~50 mV), while the shift of driving potential for Cl^−^ was smaller (from ~140 mV to ~110 mV). We expected that the increase of the “Na^+^-component” of the current would reduce Na^+^ selectivity. Indeed, the calculated P_Na+_/P_Cl−_ measured in 15 μM Ca^2+^ declined from 4.8 ± 0.6 to 2.3 ± 0.2 for wild-type TMEM16F, and from 2.1 ± 0.4 to 1.4 ± 0.1 for Q559K (Figure 5). In 1 mM Ca^2+^, switching from 15 mM NaCl to 45 mM NaCl did not significantly alter the P_Na+_/P_Cl−_ for Q559K, suggesting that Cl^−^ permeability in 15 mM NaCl was already “saturated” (Figure 5). The shift of E_rev_ by changing the monovalent ion concentration is also consistent with our proposed scenario that the shift of ion selectivity is not due to TM6 conformational change induced by Ca^2+^ entering its binding pockets, but instead, due to the change of electrostatic field along the permeation pathway.

**Figure 5.**
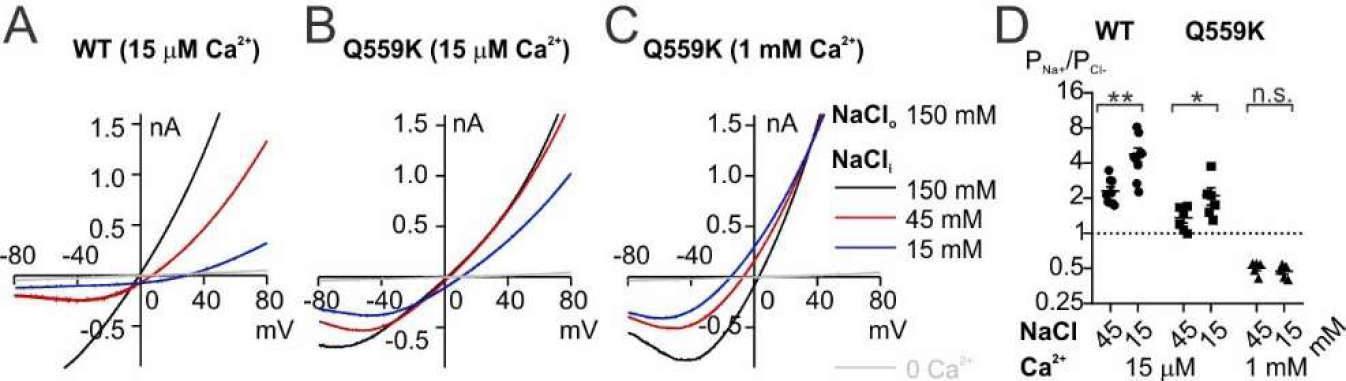
Ion selectivity is shifted with the change of intracellular NaCl concentration. (*A*) Representative I-V relationships of WT and Q559K recorded in indicated conditions. The currents were recorded with the same methods as in Figure 1E, F. Notice **<u>NOT</u>** to directly compare the shift of E_rev_ because the intracellular NaCl concentrations are varying. (*D*) Scatter plot showing the permeability ratio (P_Na+_/P_Cl−_) calculated from the shift of reversal potentials (∆E_rev_) obtained from traces as in *A*, *B* and *C*. *P* value for WT was determined with Wilcoxon test. P values for Q559K were determined with Sidak’s multiple comparisons following two-way ANOVA

### Intracellular Ca^2+^ and depolarization regulate ion selectivity synergistically

Previous studies have shown the synergy between intracellular Ca^2+^ and membrane depolarization in activating TMEM16A and TMEM16F channels (11, 18). Depolarization facilitates Ca^2+^ entry into the membrane electric field and thus reducing EC50 for channel activation. To test whether ion selectivity is also regulated synergistically by Ca^2+^ and depolarization, we held the excised membrane patch at potentials ranging from +40 to +160 mV with an increment of 40 mV (referred to as “condition potentials”) followed by a hyperpolarizing ramp from +80 mV to −80 mV (−2 V/s), to measure the reversal potentials (Figure 6A). With this experimental design, the measured E_rev_ could not be directly used to calculate the ion selectivity at the condition potentials, but the shift of values with membrane potential nonetheless provides an indication of the changing selectivity. Using this method, we measured the E_rev_ of TMEM16F Q559K channels in 15 μM, 100 μM and 1 mM Ca^2+^. Overall, currents tended to be more Cl^−^-selective as the intracellular Ca^2+^ concentration was raised. Moreover, in every given Ca^2+^ concentration, the more depolarized the condition potential, the greater the Cl^−^ selectivity (Figure 6C, E). The ability of depolarization to cause a gradual transition of ion selectivity from Na^+^ selective towards Cl^−^ selective can be attributed to intracellular Ca^2+^ being driven into the membrane electric field and/or altered driving force for monovalent ion permeation. The former is likely to have greater contribution because divalent ions have large impacts on the electrostatic field.

**Figure 6.**
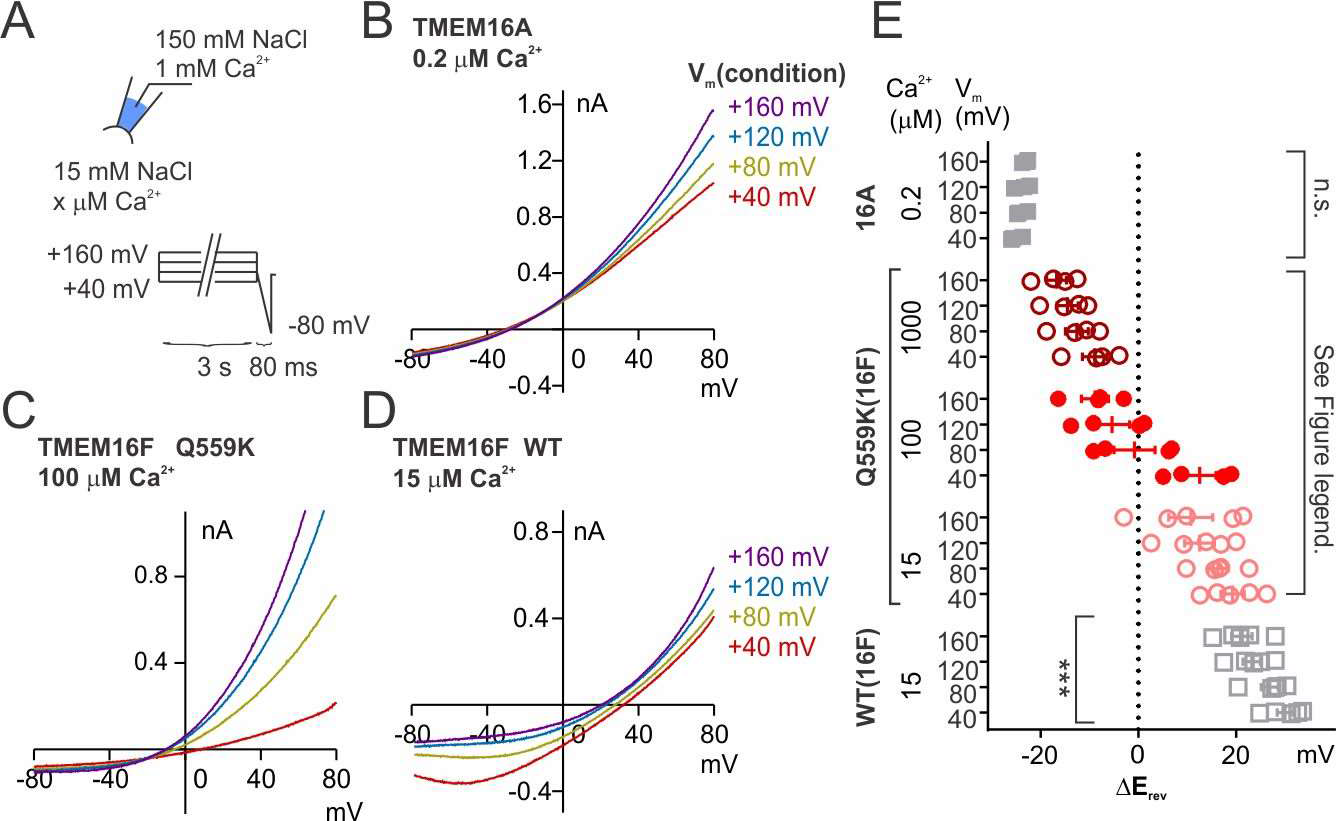
Synergy between Ca^2+^ and depolarization in shifting ion selectivity. (*A*) Recording methods. Briefly, the excised patch was held at +40 to +160 mV with an increment of 40 mV (“condition potentials”) followed by a hyperpolarizing ramp from +80 mV to −80 mV (−2 V/s), and the reversal potentials were recorded in various Ca^2+^ concentrations. (*B*~*D*) Representative I-V relationships of currents recorded in indicated conditions. (*E*) Scatter plot of the change of reversal potentials (∆E_rev_) when solution was switched from 150 mM NaCl to 15 mM NaCl. For TMEM16A and WT TMEM16F, P values were determined with one-way ANOVA. For TMEM16F Q559K, two-way ANOVA shows *P* < 0.0001 across voltages, *P* < 0.001 across Ca^2+^ concentrations, and *P* < 0.001 in their interaction

In the proposed model for TMEM16A gating (17), Ca^2+^ not only elicits conformational change of TM6, Ca^2+^ entry from the cytoplasm to occupy its binding pockets changes the electrostatic field along the permeation pathway, so as to attract Cl^−^ to the pore. In our experiment, 0.2 μM Ca^2+^ combined with a mild depolarization might have “saturated” the Cl^−^ permeability of TMEM16A channels. It is also possible that Ca^2+^ quickly retreated from the channel with the hyperpolarizing ramp, so that the E_rev_ measured with our method remained constant in spite of various condition potentials (Figure 6B, E). However, TMEM16A K584Q channels displayed reduced Cl^−^ selectivity in 0.4 μM Ca^2+^, but not in 1 mM Ca^2+^ (Figure 6–figure supplement 1). TMEM16F wild-type channels also displayed a general tendency for the ion selectivity to favor Cl^−^ as the condition potential was raised (Figure 6D, E). Given that we determined the E_rev_ with the same hyperpolarizing ramp, these measurements likely underestimated the difference of ion selectivity at different holding potentials.

### Wild-type TMEM16F ion selectivity shifts dynamically with recording conditions

The above results suggest that TMEM16F ion selectivity is shifted by the change of electrostatic field along its permeation pathway, which is determined by cytoplasmic Ca^2+^ concentration, permeating monovalent ions and membrane potential. These factors contribute to the discrepancy of TMEM16F ion selectivity reported in the literature. In whole-cell recording, the pipette solution contained 150 mM NaCl and the reversal potential was obtained with a hyperpolarizing ramp following a holding potential of +80 mV. Compared with inside-out recording as in Figure 5, higher intracellular NaCl concentration led to a larger driving force for Na^+^ permeation, and thus the channel was expected to be more selective for Cl^−^. Indeed, the shift of reversal was 1.4 ± 3.1 mV, indicative of P_Na+_/P_Cl−_ of 1.0 ± 0.1 (Figure 7A, C). With different extracellular solutions, the pattern of E_rev_ shift was seemingly indicatively of a typical Cl^−^ channel (Figure 7B, C). However, we could not calculate the selectivity of these ions due to the potential variation of P_Na+_/P_Cl−_ in these conditions.

**Figure 7.**
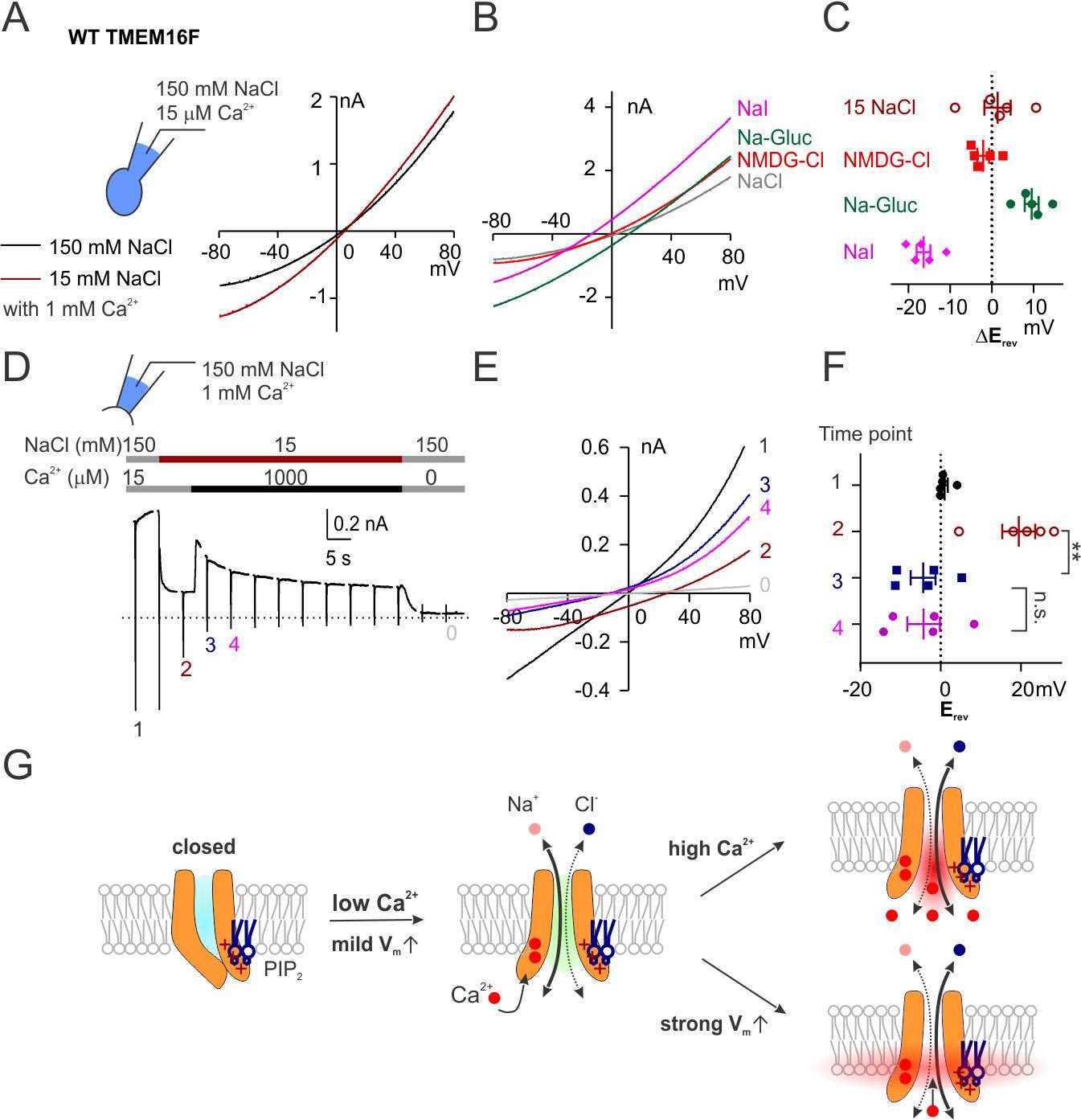
WT TMEM16F is transiently Cl^−^-selective in response to high cytoplasmic Ca^2+^. (*A*, *B*) Representative I-V relationships of wild-type TMEM16F currents recorded with whole-cell configuration in indicated bath solution. NMDG: N− methyl-D-glucamine; Gluc: gluconate. (*C*) Scatter plot of the change of reversal potentials (∆E_rev_) from recordings as in A and B. Due to the potentially varying P_Na+_/P_Cl−_, we did **<u>NOT</u>** perform statistics or use them to calculate ion permeability ratios. (D) Representative recordings of WT TMEM16F held at +80 mV with a hyperpolarizing ramp (−1 V/s) once every 5 seconds. (*E*) Representative I-V relationships of currents recorded at arrowed time points. (*F*) Scatter plot of the reversal potentials (E_rev_) at arrowed time points. P values were determined with Tukey’s multiple comparisons following one-way ANOVA. (*G*) Diagrams showing the synergy between intracellular Ca^2+^ and membrane depolarization in shifting ion selectivity via altering the electrostatic field along the permeation pathway

For most of the published results and all the data shown above, E_rev_ measurements are made after the solution change is complete and the current alteration has reached equilibrium. Here, we switched from 15 μM Ca^2+^ to 1 mM Ca^2+^ in 15 mM NaCl and immediately recorded the reversal potential with inside-out configuration. With this protocol, we found a left-shift of E_rev_ accompanied with rapid inactivation, until the current was too small to be distinguishable from endogenous current (Figure 7D, E, F). The fact that E_rev_ shifted rapidly in response to elevation of Ca^2+^ and then remained steady during the current decay supports our hypothesis that TMEM16F ion selectivity depends on Ca^2+^ concentration (when membrane potential is held constant and NaCl concentration does not change). Thus, wild-type TMEM16F dynamically undergoes the transition from being Na^+^ selective to being Cl^−^ selective with elevation of cytoplasmic Ca^2+^ concentration. This Cl^−^-selective current is transient in excised patch. In cells, the intracellular monovalent ions might be statically isotonic (~150 mM, without the ten-fold change experimentally imposed as in our excised patch recording), there might be PIP_2_ re-synthesis that could have slowed down the inactivation due to desensitization, and the summation of the “residual current” might be large enough, which could all account for the observation that TMEM16F current appears not only with a delayed activation in whole-cell recording but also with selectivity for Cl^−^ over Na^+^.

## Discussion

TMEM16F recorded with excised patch is selective for cations (6, 9). In cells with TMEM16F expression, cytoplasmic Ca^2+^ concentration increases when TMEM16F is activated (10). These findings made us wonder how TMEM16F could terminate such positive feedback of Ca^2+^ induced Ca^2+^ entry. In light of the discrepancy regarding TMEM16F ion selectivity recorded with various conditions, we set forth to test whether TMEM16F ion selectivity can be altered in response to changes of intracellular Ca^2+^. Having found that mutant Q559K channels showed minimal rundown with prolonged exposure to high cytoplasmic Ca^2+^, we determined its ion selectivity at different Ca^2+^ concentrations, revealing a shift toward Cl^−^ selectivity at high intracellular Ca^2+^. This shift of ion selectivity correlates with ion accessibility to the membrane electric field, which is determined by both ion concentration and membrane potential; divalent ions regardless of ion species are expected to exert stronger electrostatic effects than monovalent ions. This might reflect a general feature regarding a coupling of gating and permeation for channels in the TMEM16 family, and this coupling is also indicative of their intrinsic regulatory functions (Figure 7G).

Intracellular Ca^2+^ and membrane depolarization synergistically activate TMEM16F as well as Ca^2+^-activated chloride channels, TMEM16A and TMEM16B (18). In TMEM16A, membrane depolarization contributes to TMEM16A activation partly through facilitating Ca^2+^ entry into the electric field of the Ca^2+^ binding pocket (18). High concentration of intracellular Ca^2+^ allows two Ca^2+^ ions to stably bind to the binding-pocket at all tested membrane potentials, so that the channel will be stabilized at an “Ohmic” state and is constantly open regardless of the membrane potential. The amino acid sequences of Ca^2+^-binding sites are conserved across TMEM16 members, and mutation of some of these residues reduces the Ca^2+^-sensitivity (6, 9). However, TMEM16F does not have an “Ohmic” state: the channel always requires depolarization to be activated. Simply increasing cytoplasmic Ca^2+^ concentration hinders activation because the Ca^2+^ induced degradation of PIP_2_ outweighs possible further activation by Ca^2+^. Interestingly, Q559K channels show a left-shift of G-V relationship in 1 mM Ca^2+^, possibly because Ca^2+^-activation overcomes the desensitization resulting from loss of PIP_2_ (Figure 1–figure supplement 2). In addition, the depletion of PIP_2_ only desensitizes the Ca^2+^-gating of Q559K, while wild-type channel exhibits both desensitization and rundown (Figure 1A-D). One possibility is that the lysine in TM5 stabilizes the pore and prevents rundown. Alternatively, the rundown may be a secondary indicator of the shift of voltage-dependent gating. Judging from the G-V relationships, at +80 mV wild-type TMEM16F is not as fully activated in 1 mM Ca^2+^ as in 15 μM Ca^2+^ (with PIP_2_) or as the Q559K mutant in 1 mM Ca^2+^ (Figure 1–figure supplement 2). In any case, both possibilities are consistent with the model that PIP_2_ facilitates Ca^2+^-gating regulation by depolarization (11).

Recent studies of TMEM16A gating (17, 18) identify a glycine in TM6 as a hinge which allows the rearrangement of TM6 to generate the ion permeation pathway. Alanine substitution of the glycine for this hinge stabilizes the open conformation. In comparison, we found G615 in TMEM16F, the glycine at the corresponding position of G640 in TMEM16A, plays a similar role (Figure 3) (9, 10), while alanine substitution of the adjacent glycine (G614A) reduces Ca^2+^-sensitivity (Figure 3–figure supplement 1E-G). We could not find an amino acid in the lower segment of TMEM16F TM6 that plays a role similar to that of Q645 in TMEM16A, but the isoleucine at the upper segment (I612 in TMEM16F) has a function similar to that of I637 in TMEM16A (Figure 3–figure supplement 1D, G) (18), suggesting that the gating of TMEM16F ionic permeation pathway shares similar, but not identical mechanisms with TMEM16A. In TMEM16A, Ca^2+^-binding may further change the electrostatic field of the permeation pathway and reduce the energy barrier that anions need to overcome to enter the pore (17). The possibility that charges in the binding pocket and in the pore can interact electrostatically, if applicable to TMEM16F, may provide an explanation for the observation that neutralization of the lysine in the permeation pathway of TMEM16F (K641A) increases the Ca^2+^ sensitivity (10).

It is intriguing to consider the basis for Ca^2+^ regulation of TMEM16F ion selectivity. Specifically, we wonder whether cations in the permeation pathway or in the Ca^2+^-binding pocket may alter the electrostatic field of the pore. However, the fact that TM6 comprises both the gating segment and one of the pore-lining helices for the permeation pathway suggests that the physical proximity of these two “sites” makes it possible for either or both to contribute electrostatically (19, 20). Notably, in TMEM16A, introduction of negative charges to the pore can compensate for the neutralization of acidic amino acid residues in the gating pocket (17). Similarly, alteration of gating pocket charges also changes the rectification index, an indicator of ionic accessibility to the permeation pathway (17). Likewise, neutralization of basic amino acids in TMEM16F permeation pathway increases Ca^2+^-sensitivity, while neutralization of acidic residues does the opposite (9). Thus, to alter one site without influencing the other is technically challenging. Meanwhile, the recently reported TMEM16K structure has multiple Ca^2+^ ions bound to its cytoplasmic domains (21), raising the question whether there could be a reservoir in the TMEM16 cytoplasmic region to store Ca^2+^. Thus, to clearly identify the structural basis for Ca^2+^-regulation of ion selectivity might require novel technologies.

Monovalent ion concentrations in physiological conditions do not change dramatically. But the shift of ion selectivity with elevation of Ca^2+^ concentration may provide a brake to the positive feedback of TMEM16F that is activated by Ca^2+^ to mediate permeation of cations including Ca^2+^. In Figure 7, we can infer that the current at +80 mV recorded in 1 mM Ca^2+^ will be more “Cl^−^- selective” than calculated, since the hyperpolarizing ramp already leaves some time for intracellular Ca^2+^ to retreat from the membrane. The real “equilibrium point” where physiological cations and Cl^−^ are equally permeable through the TMEM16F channel, might not require intracellular Ca^2+^ as high as 1 mM and membrane potential as high as +80 mV. In a typical neuronal cell, such an ion selectivity switch might reduce the entry of Ca^2+^ into the cell and increase the permeability of Cl^−^ to drive membrane potential close to Cl^−^ equilibrium potential, both of which terminate the excitatory signal and prevent Ca^2+^-overloading. Whereas TMEM16A channels open only at depolarization when Ca^2+^ is low but display no voltage dependence in the open state when Ca^2+^ is high (18), TMEM16F might function differently with respect to its ion selectivity in response to changes in cytoplasmic Ca^2+^ concentration and membrane potential. It promotes excitation when both are low and inhibits excitation when both are high. Thus, these different members of the TMEM16 family may work as intrinsic regulatory machineries.

TMEM16F is also expressed in a variety of non-excitable cells but its function has only been investigated in a limited number of cells, such as blood cells and immune cells (1–4). Many functional studies of TMEM16F focus on lipid scrambling, and more specifically, on the exposure of phosphatidylserine to the cell surface. Here, our studies indicate that TMEM16F might also play a role in regulating membrane potential, which may regulate cellular proliferation and differentiation (22). With the broad expression pattern of TMEM16F, we envision a new perspective to examine the functions of TMEM16F.

## Materials and Methods

### Cell Culture and Molecular Biology

Mouse *Tmem16f* cDNA (sequence as in NCBI RefSeq NM_175344.4) in pmCherry-N1 vector and *Tmem16a* in pEGFP-N1 vector were generated as previously reported (11, 23). Site-directed mutagenesis was performed using standard molecular techniques with pHusion polymerase (New England Biolabs, Ipswich, MA, USA) and sequences were all verified (Quintara Biosciences, South San Francisco, CA, USA). HEK293 cells were cultured in Dulbecco’s Modified Eagle Medium (DMEM, with 4.5 g/L glucose, L− glutamine and sodium pyruvate, Mediatech, Manassas, VA, USA) containing 10% FBS (Axenia BioLogix, Dixon, CA, USA) and 1% penicillin-streptomycin, at 37°C and with 5% CO_2_. Transient transfection was performed with Lipofectamine 2000 (ThermoFisher Scientific, Waltham, MA, USA) 2 days before recording. The cDNA constructs for wild-type TMEM16F-mCherry, the mutants Q559K-, G615A-, G614A-, I612A-, G614_G615A-, N620A-, and E667Q− mCherry were stably transfected in HEK cells as previously reported (10). Briefly, the cDNAs were subcloned into pENTR1A (Addgene plasmid #17398) and transferred to pLenti CMV Hygro DEST (Addgene plasmid #17454) using Gateway cloning (24). pENTR1A no ccDB (w48-1) and pLenti CMV Hygro DEST (w117-1) were gifts from Dr. Eric Campeau and Dr. Paul Kaufman (University of Massachusetts Medical School, Worcester, MA, USA). TMEM16F-mCherry pLenti was co-transfected into HEK293FT cells with packaging plasmids pMD.2G and psPAX2, which were gifts from Didier Trono (Addgene plasmids # 12259 and #12260). Lentivirus was harvested from the transfected cells 36-48 hours post-transfection and incubated with HEK293 cells to establish stable cell lines under hygromycin selection.

### Solutions

For all electrophysiology recordings, bath solution contained 145 mM NaCl, 10 mM HEPES, 2 mM CaCl_2_, 1 mM MgCl_2_, 10 mM glucose, pH 7.2 with NaOH. For inside-out recordings, pipette solution contained 150 mM NaCl, 10 mM HEPES, 1 mM CaCl_2_, unless otherwise stated. The membrane patch was excised to form inside-out configuration in Ca^2+^-free solution: 150 mM NaCl, 10 mM HEPES, 2 mM EGTA. For solutions with Ca^2+^ < 100 μM, Ca^2+^ was added to solutions containing 2 mM EGTA or 2 mM HEDTA, and the final concentration was confirmed with Fluo-3 or Oregon Green BAPTA 5N (ThermoFisher Scientific). For whole-cell recording, the pipette solution contained 150 mM NaCl, 10 mM HEPES, 5 mM HEDTA and 4.1 mM CaCl_2_, and the final free Ca^2+^ concentration (15 μM) was confirmed with Oregon Green BAPTA 5N. The osmolality of each solution was adjusted to 290~310 mOsm/kg. For solutions with NaCl lower than 150 mM, the osmolality was balanced with addition of mannitol. All the chemicals were purchased from Sigma-Aldrich (St Louis, MO, USA).

### Electrophysiology

Cells were lifted with trypsin-EDTA (Life Technologies, Carlsbad, CA, USA) and plated onto 12 mM coverslip (Warner Instruments, Hamden, CT, USA) 3~4 days before recording. For recording, coverslips with cells were transferred to a chamber on a Nikon-TE2000 Inverted Scope (Nikon Instruments, Melville, NY, USA) and transfection was confirmed with fluorescent microscopy. For measurements of reversal potentials, a 3 M KCl salt bridge was used. Based on prediction by Clampex (Molecular Devices, Sunnyvale, CA, USA), the liquid junction potentials for 15 mM NaCl, 45 mM NaCl were −1.7 mV and −1.0 mV respectively and were only corrected for calculation of permeability ratios (P_Na+_/P_Cl-_). Patch borosilicate pipets (Sutter Instrument, Novato, CA, USA) were pulled from a Sutter P-97 puller with resistances of 2–3 MΩ for inside-out patch recordings. Solutions were puffed to the excised patch using VC3-8xP pressurized perfusion system (ALA Science, Farmingdale, NY, USA). Data were acquired using a Multiclamp 700B amplifier controlled by Clampex 10.2 via Digidata 1440A (Axon Instruments, Sunnyvale, CA, USA). All experiments were performed at room temperature (22–24°C).

### Data Analysis

All data were analyzed using pClamp10 (Molecular Devices, Sunnyvale, CA, USA), OriginLab (OriginLab Corporation, Northampton, MA, USA), and Graphpad Prism (GraphPad Software, La Jolla, CA, USA). For the measurement of Ca^2+^-sensitivity, every trace was fit to the Hill equation to generate its respective EC50 and H (Hill coefficient). The curves in the figures display the averaged current magnitudes normalized to their respective maximal values (I/Imax %). P_Na+_/P_Cl−_ values were calculated with the Goldman-Hodgkin-Katz equation:

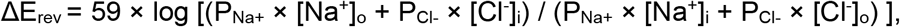

where [Na^+^]_o_, [Cl^−^]_o_, [Na^+^]_i_, and [Cl^−^]_i_ are extracellular and intracellular Na^+^ and Cl^−^ concentrations, respectively.

Significant differences were determined with Student’s t-test and ANOVA unless otherwise stated. In all cases, data represent mean ± SEM, * P < 0.05, ** P < 0.01, *** P < 0.001, **** P < 0.0001, n.s. p > 0.05.

## Acknowledgments

We thank Dr. Christian Peters and Dr. John M. Gilchrist (University of California, San Francisco) for their critical reading of the manuscript and for helpful discussions. This study is supported in part by NIH Grants R01NS069229 (to L.Y.J.), F32HD089639 (to M. H.) and by a grant from The Jane Coffin Childs Memorial Fund for Medical Research (to T.W.H.). Y.N.J. and L.Y.J. are Howard Hughes Medical Institute investigators.

## Footnotes

### Author contributions

W.Y., T.W.H., M.H. and L.Y.J. designed research; W.Y., T.W.H. and M.H. performed research; W.Y. and T.W.H. contributed new reagents/analytic tools; W.Y. analyzed data; and W.Y. wrote the manuscript and all the authors proofread and revised the manuscript.

The authors declare no conflict of interest.

---

**Figure 1–figure supplement 1.**
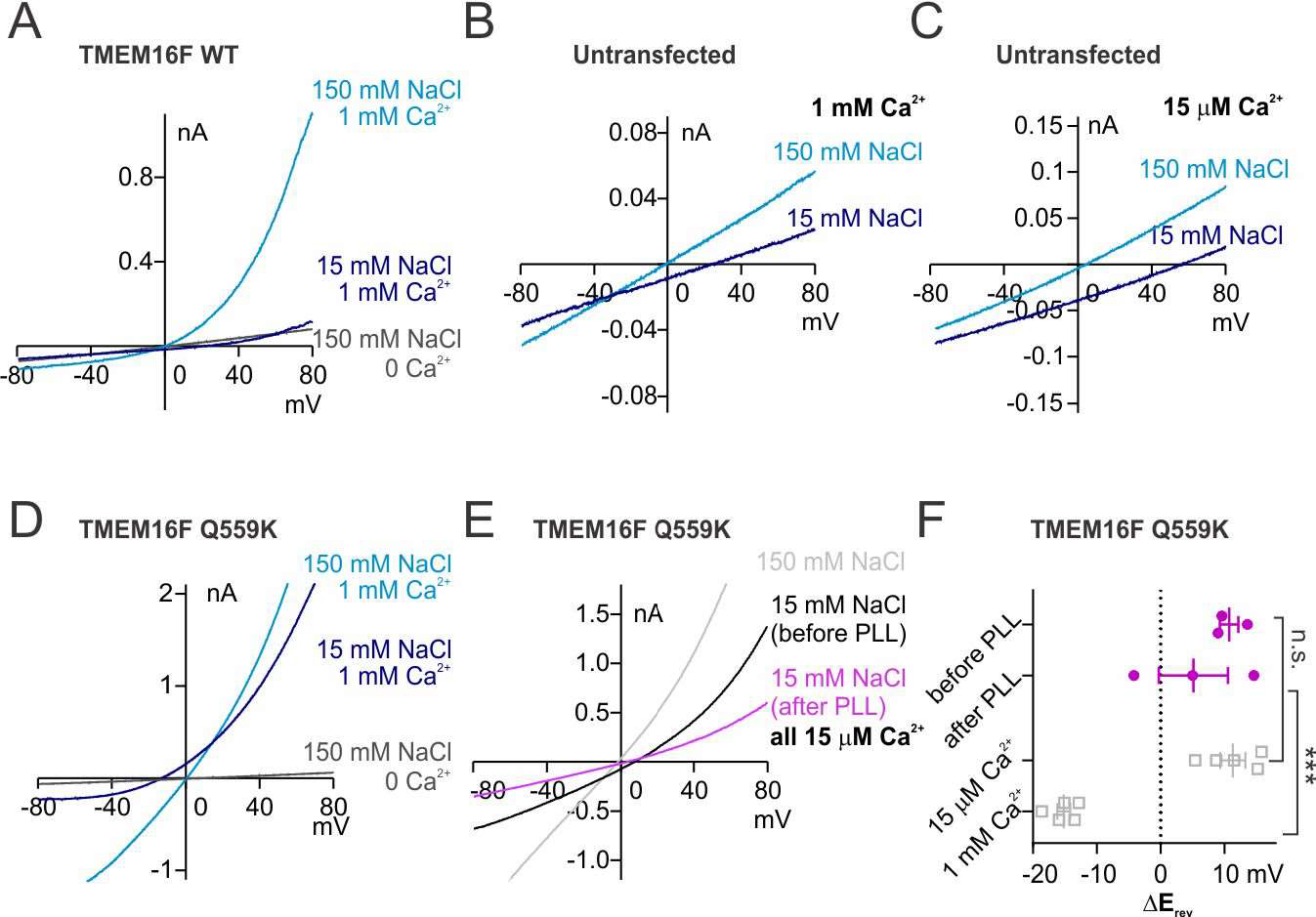
(A~D) Representative I-V relationships of currents recorded in indicated conditions. The recording protocol was the same as in Figure 1. Wild-type TMEM16F current in 1 mM Ca^2+^ is not distinguishable from HEK cell endogenous currents. (*E*) Representative I-V relationships of TMEM16F Q559K current before and after poly-L-lysine (PLL) treatment. The traces were recorded with a hyperpolarizing ramp from +80 mV to −80 mV (−1 V/s) following holding at +160 mV. (*F*) Scatter plot of the change of reversal potentials (∆E_rev_) when solution is switched from 150 mM NaCl to 15 mM NaCl. The gray values were replotted from Figure 1G. *P* values were determined with Sidak’s multiple comparisons following two-way ANOVA.

**Figure 1–figure supplement 2.**
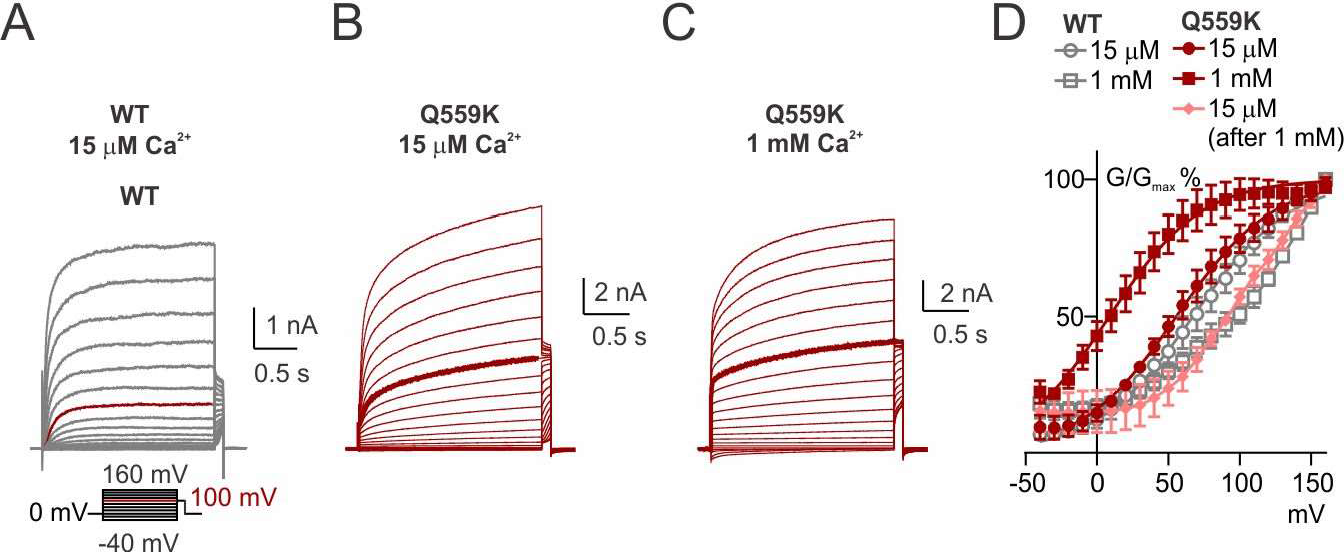
(A~C) Representative traces recorded with a voltage family protocol from −40 mV to +160 mV with 10 mV increment followed by holding at +100 mV. The traces recorded at +100 mV are highlighted for comparison. (D) Averaged G-V relationships of the currents recorded in indicated conditions. Tail current (at +100 mV) magnitudes were measured from traces as in A~C. For WT and Q559K in 15 μM Ca^2+^ and Q559K in 1 mM Ca^2+^, currents of each cell were fitted to sigmoidal equation and the current magnitudes were normalized to their respective maximal values. For WT in 1 mM Ca^2+^ and Q559K after 1 mM Ca^2+^ treatment, the current magnitudes were directly normalized to their values at +160 mV.

**Figure 1–figure supplement 3.**
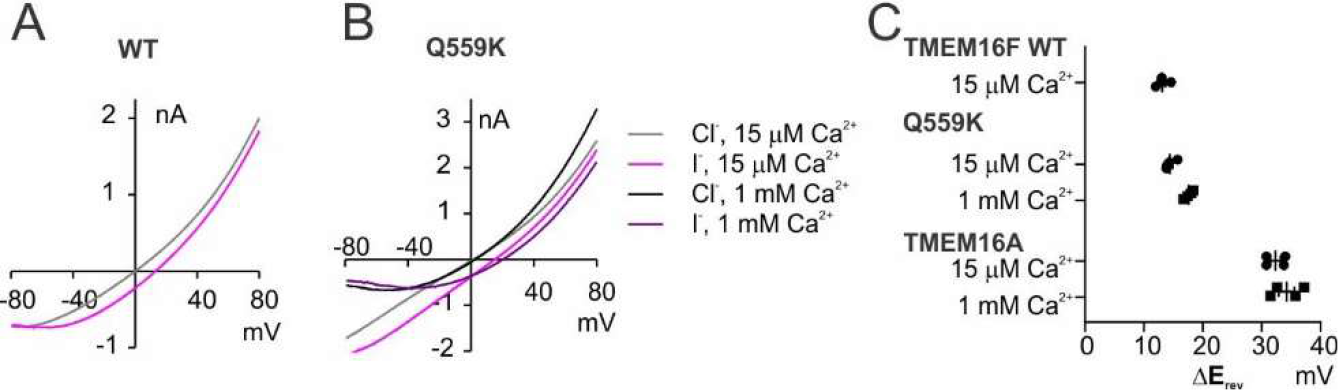
(A, B) Representative I-V relationships of currents recorded in indicated conditions showing the shift of reversal potentials when the bath (intracellular) solution was switched from 150 mM NaCl to 150 mM NaI. (C) Scatter plot of the change of reversal potentials (∆E_rev_) when solution was switched from 150 mM NaCl to 150 mM NaI. Because the interpretation methods for bi-ionic condition were not applicable here, we did **<u>NOT</u>** perform statistics and instead showed the values directly.

**Figure 3–figure supplement 1.**
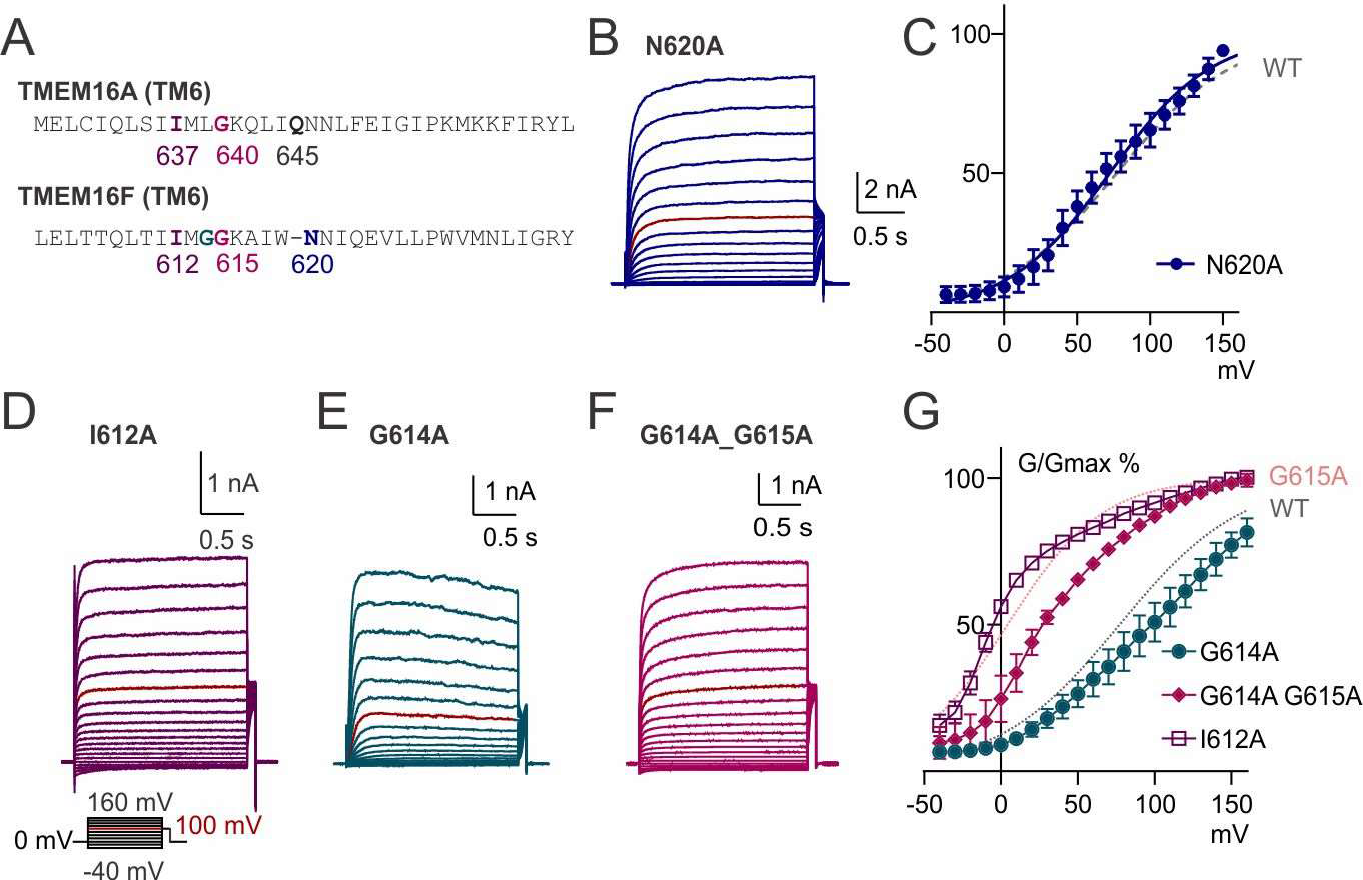
(*A*) Alignment of TM6 sequences of TMEM16A and TMEM16F. The numbering for TMEM16A represents the isoform as used by Peters, C et al. (18). (*B*, *D*~*F*). Representative traces of TMEM16F mutants recorded in 15 μM Ca^2+^ with a voltage family protocol as in Figure 1_Supplement 2. Currents recorded at +100 mV are highlighted for comparison. (*C*, *G*) Averaged G-V relationships of indicated TMEM16F mutants. The method for data analysis was the same as that for WT in 15 μM Ca^2+^. The traces for WT and G615A were replotted from Figure 1_Supplement 2D and Figure 3B.

**Figure 4–figure supplement 1.**
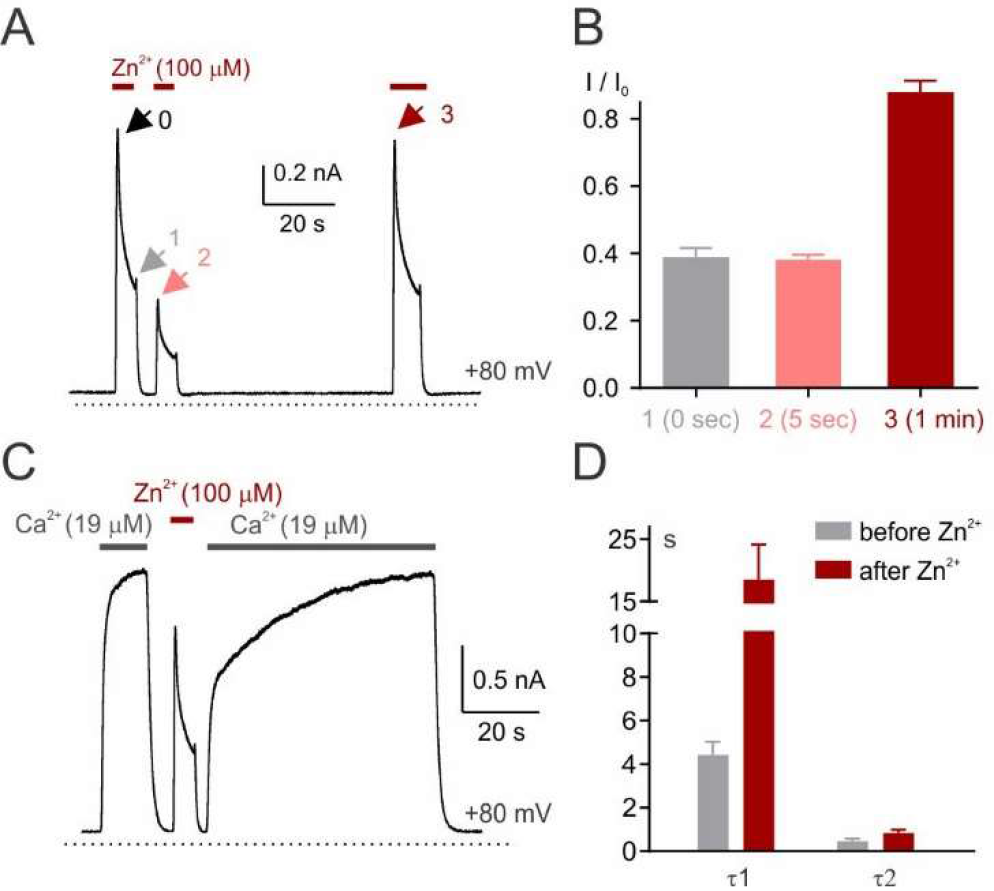
(*A*) Representative recordings of WT TMEM16F current in response to multiple applications of Zn^2+^. (*B*) Averaged current magnitudes measured at arrowed time points normalized to the respective initial magnitudes (I0), indicating that Zn^2+^-inactivation is reversible, not due to PIP_2_ depletion. (*C*) Representative recordings of WT TMEM16F current showing Ca^2+^ activation is not desensitized by Zn^2+^-treatment, although the activation is slower. (*D*) Averaged time constants for Ca^2+^-activation. The time constants were obtained by fitting the activation trace with a two-term exponential curve.

**Figure 6–figure supplement 1.**
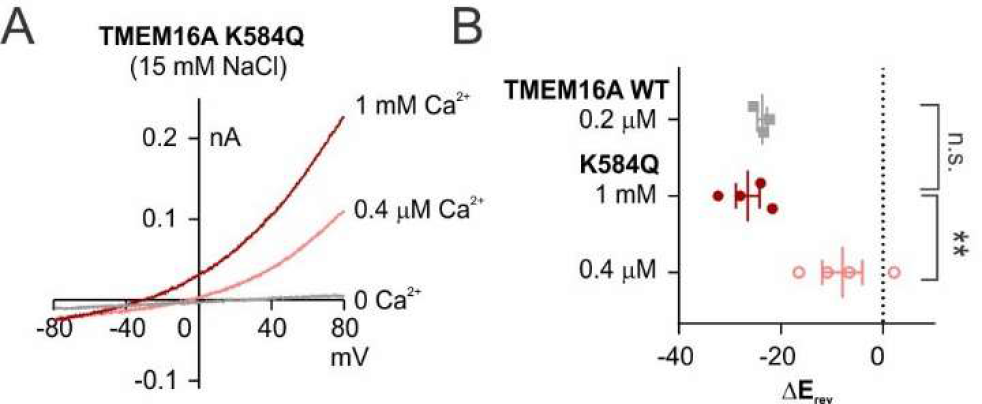
(*A*) Representative I-V relationships of TMEM16A K584Q currents recorded in indicated concentrations of Ca^2+^ in 15 mM NaCl bath solution. (*B*) Scatter plot of the change of reversal potentials (∆E_rev_) when solution was switched from 150 mM NaCl to 15 mM NaCl. P values were determined with Sidak’s multiple comparisons following one-way ANOVA.

